# *Tceal7* is a BRG1-regulated target of calcineurin signaling that promotes myoblast differentiation

**DOI:** 10.64898/2026.01.20.700469

**Authors:** Sandhya Yadav, Tapan Sharma, Teresita Padilla-Benavides, Anthony N. Imbalzano

## Abstract

The transcriptional control of skeletal muscle differentiation requires the coordinated activity of lineage-defining transcription factors, signal-responsive regulators, chromatin modifiers, and ATP-dependent chromatin remodeling enzymes. Here, we identify TCEAL7, a member of the X-linked, poorly characterized TCEAL family of proteins, as a direct downstream target of BRG1-containing mammalian SWI/SNF (mSWI/SNF) complexes and calcineurin signaling during myoblast differentiation.

Analyses of previously published datasets showed that pharmacological inhibition of mSWI/SNF bromodomains or knockdown of the BRG1 ATPase, but not knockdown of the homologue BRM ATPase, significantly reduced *Tceal7* expression in differentiating C2C12 myoblasts. We demonstrate that BRG1 occupancy at the *Tceal7* promoter increased during differentiation, paralleling the induction of *Tceal7* expression and nuclear accumulation of TCEAL7 protein. BRG1 functions in part by integrating calcium-dependent cues via the phosphatase calcineurin (Cn); we also determined that Cn knockdown or pharmacological inhibition of Cn suppressed *Tceal7* expression and impaired myoblast differentiation. The data suggest that both BRG1-driven chromatin remodeling and Cn signaling converge on *Tceal7* regulation. Functionally, *Tceal7* knockdown altered cell proliferation and disrupted myoblast differentiation, at least in part due to reduced expression of *Myogenin*, which encodes a transcription factor that is an essential differentiation determinant. RNA-seq analysis revealed broad dysregulation of myogenic, metabolic, and cell-cycle gene programs in *Tceal7*-deficient cells, including changes in cyclin-dependent kinase-regulated pathways consistent with prior reports linking TCEAL7 to cell-cycle control. Together, these findings identify TCEAL7 as a necessary component of the myogenic regulatory network whose expression is controlled by BRG1-dependent chromatin remodeling and Cn activity.

## INTRODUCTION

The TCEAL family of proteins is encoded by a cluster of X chromosome-linked genes that evolutionarily conserved in mammals [1, 2]. The first member of the family identified, now called TCEAL1 [3], was reported to share homology with the transcription elongation factor A (TCEA), also known as TFIIS [4–7]. Specifically, TCEAL1 shared limited homology with the amino acid sequences identified as interacting with RNA polymerase II and was reported to have a zinc (Zn) finger-“like” domain resembling the Zn finger existing in TFIIS [3]. Other TCEAL proteins and the genes encoding them were identified and named based on their homology to TCEAL1. There are nine TCEAL proteins in humans and seven in mice [2]. The TCEAL family members are presumed to play a role in transcription elongation or, since TFIIS was later shown to also be involved in transcription initiation by RNA polymerase II [5, 7], transcription regulation in general. However, to our knowledge, there are no reports demonstrating that any of the TCEAL proteins bind Zn, bind RNA polymerase II, or function directly in transcription. We recently compared the known structure of TFIIS to the structures of each TCEAL protein as predicted by Alphafold2 [8, 9] and proposed that it is likely that none of the TCEAL proteins bind either Zn or RNA polymerase II [10]. Functionally, TCEAL protein and/or mRNA levels frequently have been identified as mis-regulated, both positively and negatively, in many types of cancer [10]. Several TCEAL proteins have links to protein de-ubiquitination [11–13], and most are predicted to be phosphoproteins [10]. However, a unifying function, if one exists, remains unknown.

Mammalian SWI/SNF (mSWI/SNF) enzymes are ATP-dependent chromatin remodelers [14–16]. These enzymes are multi-subunit complexes with significant diversity in assembly, resulting in the formation of many different enzyme complexes [17, 18]. However, biochemical studies of enzyme assembly revealed that there are three main sub-families, each characterized by unique subunits [19]. The enzymes alter nucleosome structure to facilitate access to DNA during transcription, replication, recombination and repair [20–23]. Consequently, the mSWI/SNF enzymes are critical for development and for many normal cellular processes. Not surprisingly, mis-regulation or mutation/deletion of different subunits are associated with many types of cancer [24–26], and it has been reported that ∼20% of all human cancers contain mis-regulated or mutated mSWI/SNF enzyme subunits [25].

mSWI/SNF chromatin remodelers have been shown to be critical for the differentiation of myoblasts in culture and during adult skeletal muscle differentiation and function [27–33]. Altered expression or mutation of mSWI/SNF enzyme subunits results in mis-regulation of genes essential for cell cycle regulation proteins [34, 35], differentiation [27, 36, 37], and those of the Wnt signaling pathway [38–40], which is critical for myogenesis [41, 42].

Our group has reported the deleterious effects of a drug called PFI-3 on myoblast differentiation [32]. PFI-3 is a bromodomain inhibitor that specifically binds to bromodomains in three distinct mSWI/SNF subunits (BAF180 and the mutually exclusive ATPases, BRM and BRG1), but no other proteins [43, 44]. We have also examined the effects of siRNA treatment targeting BRG1, BRM or both [45]. PFI-3 treatment and knockdown of BRG1, BRM or both inhibited myoblast differentiation in culture and PFI-3 inhibited in adult myogenesis after injury *in vivo* [32, 45]. Among the genes that were sensitive to PFI-3 treatment and BRG1, but not BRM, siRNA was a TCEAL gene, *Tceal7*.

TCEAL7 was of interest because it has been implicated in myogenesis [46–50]. The *Tceal7* gene was differentially expressed during skeletal muscle regeneration following injury in adult mice, while *Tceal7* mRNA was expressed only in skeletal muscle during mouse embryogenesis [47]. *Tceal7* gene expression changes were also identified in skeletal muscle using different mouse disease models [50, 51] and during *in vitro* mouse myoblast differentiation [52]. Studies of the regulation of *Tceal7* gene expression in skeletal muscle originally utilized transgenic mice to identify a 0.7 kb portion of the sequences upstream of the *Tceal7* transcription start site (TSS) that could recapitulate expression in developing embryonic skeletal muscle [47]. Mutation of the E boxes in this sequence, which are binding sites for the myogenic regulatory factor (MRF) family of basic helix-loop-helix transcription factors that includes MyoD1, myogenin, Mrf4, and Myf5 [53], resulted in loss or reduction of transgene expression [47, 49]. Each member of the MRF family stimulated *Tceal7* expression and at least one of the MRFs, MyoD1, could bind to the *Tceal7* E boxes [47]. Subsequent work showed that additional transcriptional regulators known to impact myogenic gene expression, MEF2C and CREB1, could cooperate with MyoD1 to regulate *Tceal7* expression [49].

Functionally, myoblasts overexpressing *Tceal7* showed enhanced differentiation capacity. The expression of several cell cycle regulators was surveyed; the only tested mRNA to show a change was *Cdkn1b*, which encodes p27 and which was elevated [48]. p27 is a cyclin dependent kinase inhibitor that functions to reduce cell proliferation [54], which likely at least partially explains the effect of *Tceal7* overexpression. Transgenic mice expressing *Tceal7* in skeletal muscle after the initiation of skeletal muscle development did not impact embryogenesis or post-natal growth for the first three weeks [54]. However, the overall size and body weight was altered subsequently, at least in part due to reduced cross-sectional area of examined skeletal muscles [54]. Additionally, direct interaction between TCEAL7 and CDK1 was observed [54]. CDK1 is a cyclin-dependent kinase that, when coupled with a partner cyclin, phosphorylates target proteins involved in protein synthesis [55]. The authors proposed that the TCEAL7 protein repressed CDK1 activity, thereby negatively impacting protein synthesis and affecting myofiber growth [54]. TCEAL7 may therefore impact both myoblast differentiation and myofiber growth. Another study showed that knockdown of either *Tceal7* or *Tceal5* did not affect differentiation in culture but simultaneous knockdown of both did [46].

Here, we show that *Tceal7* expression is controlled by BRG1-containing SWI/SNF chromatin-remodeling complexes. *Tceal7* is robustly induced during myogenic differentiation, localizes to the nucleus, and is essential for proper *Myogenin* expression, myotube fusion, and activation of myogenic transcriptional networks. shRNA-mediated *Tceal7* knockdown severely impaired myoblast differentiation and broadly reduced the expression of myogenic genes, while inducing signaling pathways that likely hinder, rather than support, normal myoblast differentiation. Together, our findings support a model where *Tceal7* is a critical regulator of the transcriptional and signaling programs that drive myoblast differentiation.

## MATERIALS AND METHODS

### Antibodies

The primary antibodies against BRG1 that were used for western blot and chromatin IP (sc-17796; 1:1,000 and sc-17796 G-7X, 3-4 μL per μg of chromatin, respectively), against vinculin (sc-25336; 1:1000), and against Lamin β1 (sc-56144; 1:10,000) were purchased from Santa Cruz Biotechnologies. Myosin Heavy Chain monoclonal antibody (MF20, deposited by D. A. Fischman) was purchased from the Developmental Studies Hybridoma Bank, University of Iowa. The calcineurin antibody was from Cell Signaling Technology, Inc (2614; 1:1,000). The secondary antibodies used were goat anti-rabbit and -mouse coupled to HRP (31460, 31430 respectively; 1:5,000) and were acquired from Thermo Fisher Scientific.

### Mammalian cell culture

C2C12 cells are a karyotypically abnormal subclone of a spontaneously immortalized mouse skeletal myoblast line widely used to model myoblast proliferation and differentiation [56]. C2C12 cells were obtained from ATCC (Manassas, VA) and maintained at sub-confluent densities in proliferation medium consisting of Dulbecco’s Modified Eagle Medium (DMEM; 11965118, ThermoFisher Scientific) supplemented with 10% fetal bovine serum (FBS) and 1% penicillin-streptomycin in a humidified incubator at 37°C with 5% CO_2_. C2C12 differentiation was initiated when cultures reached approximately 80% confluence. Cells were switched to differentiation medium containing DMEM supplemented with 2% horse serum, 1% insulin-transferrin-selenium A (Invitrogen), and 1% penicillin-streptomycin. Differentiated myoblasts were collected at the time points indicated in the figure legends and processed for further analyses. PFI-3 (15267; Cayman Chemicals) was added to cell culture media at a final concentration of 50 μM at the time of induction of differentiation [32].

### siRNA transfection of C2C12 myoblasts

The following siRNA oligos were purchased from Dharmacon Horizon Discovery Ltd., United States. siRNA against *Brg1* (siGENOME mouse Smarca4 pool #M-041135-01-0020, 50nM; siGENOME mouse *Smarca4* #D-041135-03-0050, 25nM, referred to as si*Brg1*-A; and siGENOME mouse *Smarca4* #D-041135-04-0050, 25nM, referred to as si*Brg1*-B). The non-targeting siRNA (SMARTpool ON-TARGETplus scrambled # D-001810-10-20, 50 nM) was used as control. C2C12 cells were transfected as previously described [32] using indicated concentrations of different siRNAs and harvested at indicated times for further analysis

### shRNA-mediated gene knockdown

shRNA-mediated knockdown was done using shRNAs and procedures as previously described [57].

### Western blot analysis

C2C12 cell samples from three independent experiments were washed twice with PBS and scraped into 1 mL PBS using a cell lifter. Cells were pelleted by centrifugation and lysed in 500 μL RIPA buffer (50 mM Tris–HCl, pH 7.4; 150 mM NaCl; 1 mM EDTA; 1% NP-40; 0.25% sodium deoxycholate) supplemented with protease inhibitor cocktail (P8340, Sigma Aldrich). Lysates were sonicated in an ultrasonic bath for three 30 s on/off cycles at 4°C. Samples were centrifuged at 14,000 × *g* for 10 min at 4°C, and supernatants were collected. Protein concentrations were measured using the Pierce™ BCA assay (ThermoFisher Scientific). A total of 20 μg protein mixed with loading dye (5% β-mercaptoethanol, 0.02% bromophenol blue, 30% glycerol, 10% SDS, 250 mM Tris-Cl, pH 6.8) was boiled at 95°C for 10 min and resolved on 10% SDS-PAGE gels, followed by transfer to Immobilon-P PVDF membranes (Merck Millipore). Membranes were blocked in 5% non-fat milk for 30 min and incubated overnight at 4°C with primary antibodies diluted in 2% milk in PBS or TBS. After three 5-min washes in TBS with 0.1% Tween-20, membranes were incubated for 1 h at room temperature with HRP-conjugated species-specific secondary antibodies, followed by three additional washes. Bands were detected by chemiluminescence using ECL Plus (GE Healthcare) on an Amersham Imager 600. Band intensities from three independent experiments were quantified using ImageJ [58].

### Cell proliferation assays

C2C12 myoblasts were seeded at 1 × 10^4^ cells/cm^2^, and samples were collected at 24, 48, and 72 h post-plating. Cells were trypsinized, washed three times with PBS, and counted using a Cellometer Spectrum (Nexcelom Biosciences).

### RT-qPCR Gene Expression Analysis

Total RNA was isolated from three independent biological replicates of differentiated C2C12 myoblasts using TRIzol (Invitrogen) according to the manufacturer’s instructions. cDNA was synthesized from 500 ng RNA using random primers and SuperScript III reverse transcriptase (Invitrogen). Quantitative PCR was performed with Fast SYBR Green 2X Master Mix (Applied Biosystems), and final reaction volume was adjusted to 10 μL per reaction. All qPCRs were run in QuantStudio 3 RT-PCR machine (Applied Biosystems) using primers listed in **Supplemental Table 1**. ΔC_T_ was calculated as the C_T_ of the target gene minus the C_T_ of the housekeeping gene (*Eef1a1*). ΔΔC_T_ was then obtained by subtracting the average ΔC_T_ of the control group from the ΔC_T_ of each sample. Relative expression was determined using the 2^⁻ΔΔCT^ method [59].

### Immunocytochemistry Analyses

C2C12 myoblasts (control and knockdown) were fixed overnight in 10% formalin-PBS at 4°C, washed with PBS, and permeabilized for 10 min in PBS with 0.2% Triton X-100. Immunocytochemistry was performed using the myosin heavy chain hybridoma supernatants. Staining was developed using the Universal ABC kit (PK-6200; Vector Laboratories, United States) and HRP DAB substrate (SK-4100; Vector Laboratories, United States) according to the manufacturer’s instructions.

### Chromatin Immunoprecipitation (ChIP) Assays

ChIP assays were performed as previously described [57]. Briefly, differentiating C2C12 myoblasts were cross-linked with 1% formaldehyde for 10 min at room temperature and quenched with 125 mM glycine for 5 min. Cells were washed twice with ice-cold PBS containing protease inhibitors and lysed in 1 mL of ice-cold SimpleChIP ® Enzymatic Cell Lysis Buffer A (14282; Cell Signaling Technology). Nuclei were pelleted at 3000 × g, washed in buffer B (14231; Cell Signaling Technology) as recommended by the manufacturer. Chromatin was digested with 1000 U micrococcal nuclease (M0247S; New England Biolabs) for 30 min at 37°C. Reactions were stopped with 5 μL of 0.5 M EDTA. Nuclei were pelleted, resuspended in 400 μL ChIP buffer (SimpleChIP ® Chromatin IP Buffers, Cell Signaling Technology) supplemented with protease inhibitors, sonicated for 10 min (30 s on/30 s off, medium intensity) using a Bioruptor UCD-200 (Diagenode), and centrifuged at 21,000 × g for 5 min. Fragmented chromatin (200-500 bp) was confirmed by agarose gel electrophoresis.

Chromatin was incubated for 2 h at 4°C with antibodies against BRG1 and with an anti-IgG antibody as a negative control was included. Immunocomplexes were captured with 20 μL Dynabeads (Thermo Fisher Scientific) after overnight incubation at 4°C, washed three times with low-salt ChIP buffer and once with high-salt buffer, and eluted in 100 μL elution buffer (0.1 M NaHCO_3_, 1% SDS) for 30 min at 65°C. Samples were treated with RNase (1 μL, 0.5 mg/mL, 30 min, 37°C), then reverse-cross-linked overnight at 65°C with 6 μL 5 M NaCl and 1 μL proteinase K (1 mg/mL). DNA was purified using the ChIP DNA Clean & Concentrator kit (Zymo Research). Enriched DNA was analyzed by qPCR using SYBR Green master mix and quantified by the 2^(ΔCT sample – ΔCT IgG)^ method. The data are shown relative to the results determined for IgG controls. Primer sequences are listed above.

### RNA-sequencing analysis

Total RNA (duplicate samples per condition) was assessed for quality and concentration using the UMass Chan MBCL Fragment Analyzer. Samples with RIN ≥ 7 and 28S/18S ≥ 1.0 were submitted to BGI Americas for library preparation and RNA-seq. After adapter trimming, clean reads were aligned to the *mm10* reference transcriptome using HISAT2, and gene expression was quantified with featureCounts [60]. Differentially expressed genes were identified using DESeq2 [61], applying thresholds of log_2_ fold change ≥ ±0.5 and adjusted P < 0.05. Dot plots were generated in RStudio using ClusterProfiler [62]. Pathway enrichment was performed with the PANTHER database [63, 64].

### Statistical Analysis

Statistical analyses were performed using GraphPad Prism 7.0b. Student’s t-test was used to compare the differences between the two groups. P values < 0.05 were considered statistically significant.

## RESULTS

### BRG1 is required for *Tceal7* expression during myoblast differentiation

We previously performed RNA-seq on differentiating C2C12 cells that had been treated with PFI-3, a bromodomain inhibitor that binds to and interferes with bromodomains on three mSWI/SNF subunits: the mutually exclusive ATPases, BRG1 and BRM, and the BAF180 subunit specific to the PBAF subfamily of mSWI/SNF complexes [32, 44]. In that study, we demonstrated that the effects of PFI-3 were due to inhibition of BRG1 and BRM; BAF180 was dispensable for myoblast differentiation [32]. We also previously analyzed differentiating C2C12 cells treated with siRNA targeting *Brg1*, *Brm*, or both [45]. We noted that *Tceal7*, a gene previously implicated in skeletal muscle development [46–50], was down regulated in differentiating C2C12 cells treated with PFI-3 and siRNA targeting *Brg1*, but not in cells treated with siRNA targeting *Brm* (**Fig. 1A**). To directly validate *Tceal7* as a BRG1 target, we performed siRNA-mediated knockdown experiments using these cells. *Brg1* knockdown markedly reduced *Tceal7* mRNA and TCEAL7 protein abundance (**Fig. 1B,C**). These findings demonstrate that *Tceal7* is transcriptionally controlled by BRG1 and functions as a downstream effector within BRG1-based mSWI/SNF-mediated myogenic regulatory networks.

**Figure 1.**
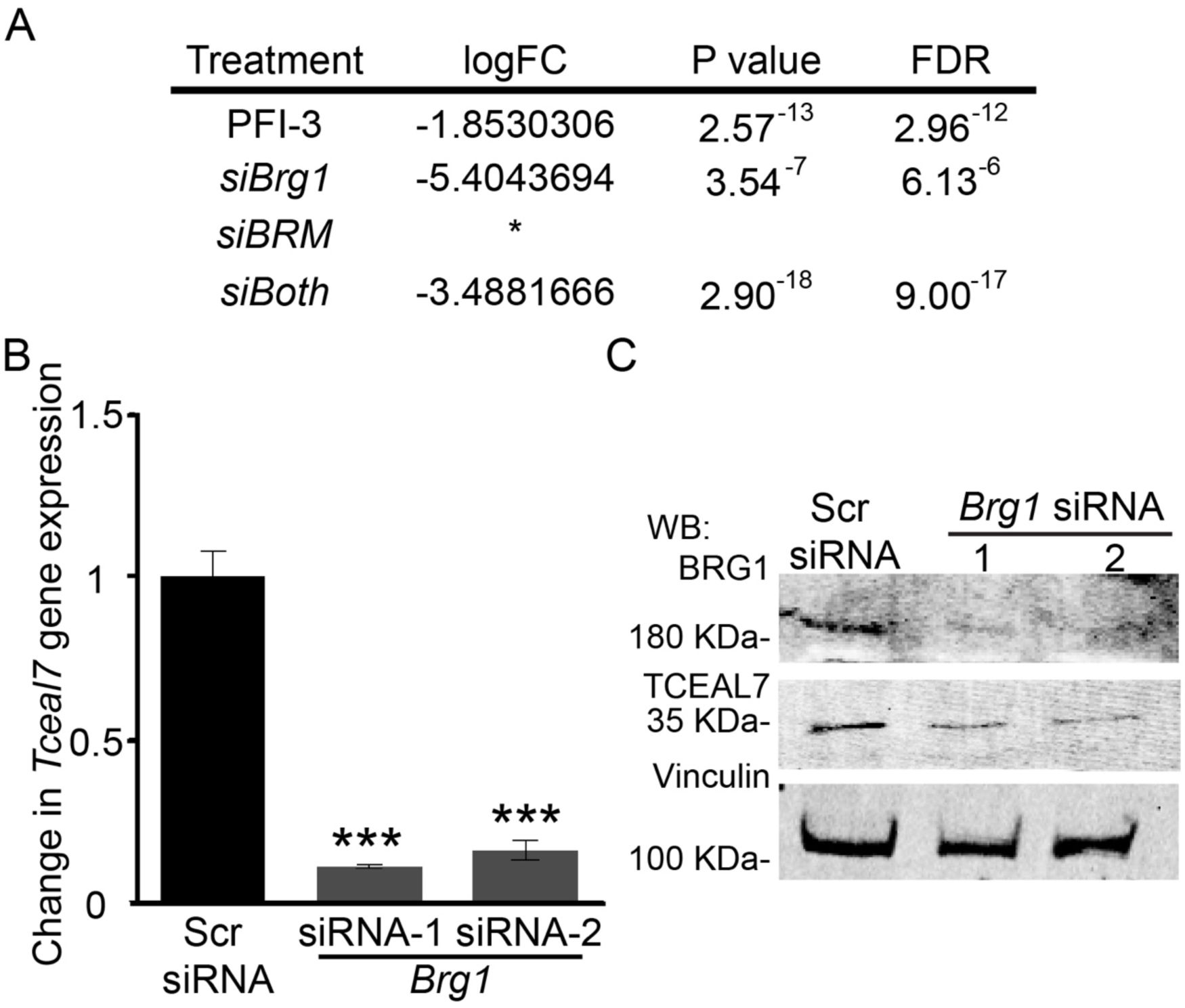
*Tceal7* expression is regulated by BRG1 but not BRM. **(A)** *Tceal7* gene expression levels in differentiating C2C12 myoblasts treated with PFI-3 (from GSE151218) or treated with siRNAs targeting *Brg1* or *Brm* (from GSE196283). The asterisk indicates the gene was not differentially expressed. (**B**) Steady state *Tceal7* mRNA expression in C2C12 cells transfected with control scrambled sequence (Scr) or one of two different *Brg1* siRNAs, normalized to *Eef1a* expression. **(C)** Western blots showing reduced TCEAL7 protein levels upon *Brg1* knockdown. Vinculin served as the loading control. Data represents 3 independent biological experiments ± SD. *P < 0.05; ***P < 0.001.

### *Tceal7* is induced during C2C12 myoblast differentiation and its promoter is inducibly bound by BRG1 during differentiation

To determine how *Tceal7* expression changes during C2C12 myoblast differentiation, we examined expression at 24 hour intervals from the onset of differentiation to 72 hours post-differentiation. Immunohistochemistry of the differentiation marker myosin heavy chain (MHC) showed the transition from single C2C12 myoblasts to elongated, muti-nucleated myotubes (**Fig. 2A**), thereby providing a visual assessment of the state of the cells at each timepoint. *Tceal7* transcript levels increased during this period (**Fig. 2B**), and immunoblotting revealed a similar increase in the protein levels of TCEAL7 and myogenin, an early myogenic marker (**Fig. 2C**). The increase in TCEAL7 protein level is more rapid than is the increase in *Tceal7* mRNA. This suggests the possibility of post-transcriptional regulation.

**Figure 2.**
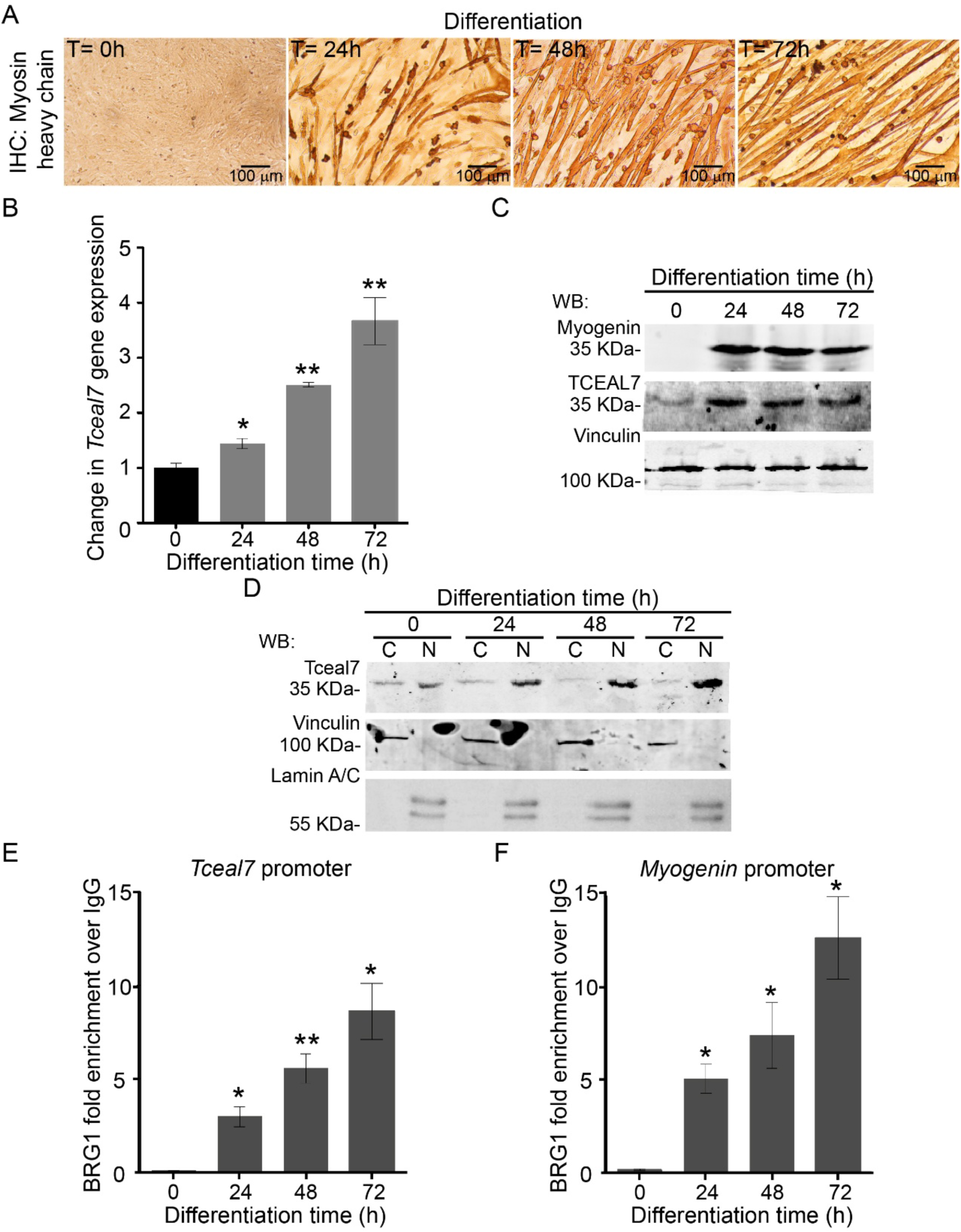
TCEAL7 is induced during C2C12 differentiation and accumulates in the nucleus and the cytoplasm; the *Tceal7* promoter is bound by BRG1. **(A)** Representative light microscopy images of myosin heavy chain staining of differentiating myoblasts at 0, 24, 48, and 72 h. **(B)** Steady state levels of *Tceal7* mRNA levels during the time course of differentiation, normalized to *Eef1a*. **(C)** Representative western blot showing TCEAL7 and myogenin protein expression over the differentiation time course; vinculin was used as the loading control. **(D)** Representative western blot analysis of TCEAL7 in cytoplasmic (C) and nuclear (N) fractions at indicated time points. To determine purity of fractions, vinculin was used as the cytoplasmic extract control and Lamin A/C was used as the nuclear extract control. (**E-F)** ChIP analysis showing BRG1 binding at the *Tceal7* (E) and *Myogenin* (F) promoters across the differentiation time course. Values were normalized to values obtained for an IgG ChIP. Data represent 3 independent biological experiments ± SD. *P < 0.05; **P < 0.01.

### TCEAL7 is found in both the cytoplasm and the nucleus of differentiating myoblasts

Evidence linking TCEAL7 to CDK1 function [48] and inhibition of the ability of the transcription factor NF-KB to bind DNA [65] suggest TCEAL7 is a nuclear protein, and HA-tagged TCEAL7 localizes to the nucleus [66]. Furthermore, analysis of human samples showed that TCEAL7 was present in the nucleus of normal gastric tissues as well as in differentiated gastric tumors [67].

To determine the subcellular distribution of TCEAL7 in differentiating myoblasts, we analyzed cytoplasmic and nuclear fractions as a function of time of differentiation. TCEAL7 was detected in both the nucleus and the cytoplasm at the onset of differentiation but became enriched in the nuclear fraction with increasing differentiation time (**Fig. 2D**). The data reinforce the idea that TCEAL7 is found in the nucleus but leave open the possibility of both cytoplasmic and nuclear functions for TCEAL7.

### The *Tceal7* promoter is inducibly bound by BRG1 during differentiation

Work from others has demonstrated that myogenic regulatory factors (MRFs) activate *Tceal7* expression via E box elements, which are MRF binding sites, in the *Tceal7* promoter [47, 48]. As BRG1-based mSWI/SNF complexes cooperate with MRFs to activate myogenic gene expression [27, 28, 68], we predicted that BRG1 should be bound to the *Tceal7* promoter when it is activated. BRG1 ChIP assays showed inducible binding to the *Tceal7* promoter that increased as a function of time of differentiation (**Fig. 2E**). BRG1 binding correlated with the increase in mRNA as well as with the timing of BRG1 binding to the *myogenin* promoter, a known BRG1 target ([27, 28, 68]; (**Fig. 2F**). Together, these data establish *Tceal7* as a differentiation-induced gene that is likely directly regulated by BRG1.

### Calcineurin A regulates *Tceal7* expression and myoblast fusion

Since BRG1 cooperates with calcineurin (Cn) to activate myogenic gene expression and promote myoblast differentiation [57, 69], we next tested whether this phosphatase also contributes to the regulation of *Tceal7*. We generated myoblasts using shRNA targeting *CnA* (the A subunit of the Cn protein; [70]) and used immunohistochemistry analyses of differentiating myoblasts stained against myosin heavy chain to confirm that *CnA* knockdown disrupted myotube formation, producing thinner and poorly fused myotubes relative to controls (**Fig. 3A**). Q-PCR analysis of these cells confirmed knockdown of *CnA* (**Fig. 3B**). *CnA* knockdown reduced the levels of *Tceal7* mRNA (**Fig. 3C**). Immunoblotting showed that CnA and TCEAL7 protein levels (**Fig. 3D**), were also reduced. Together, these results show that Cn signaling is required for proper activation of *Tceal7* during myoblast differentiation. To demonstrate that CnA activity modulates TCEAL7 protein abundance, we treated differentiating C2C12 cells with the Cn inhibitor FK506 [71]. TCEAL7 protein levels decreased across differentiation time points in FK506-treated cultures (**Fig. 3E**), confirming that Cn activity is necessary to sustain *Tceal7* expression. Because BRG1 function is sensitive to CnA activity [69], we examined the binding of BRG1 to the *Tceal7* promoter as well as to the promoter driving the expression of the myogenic marker gene, *Myogenin*, which we have previously shown is sensitive to *CnA* inhibition [69]. BRG1 binding to both promoters was reduced in the presence of shRNA targeting CnA (**Fig. 3F**). These results reveal that Cn signaling promotes *Tceal7* expression at least in part by facilitating BRG1 recruitment to the *Tceal7* promoter.

**Figure 3.**
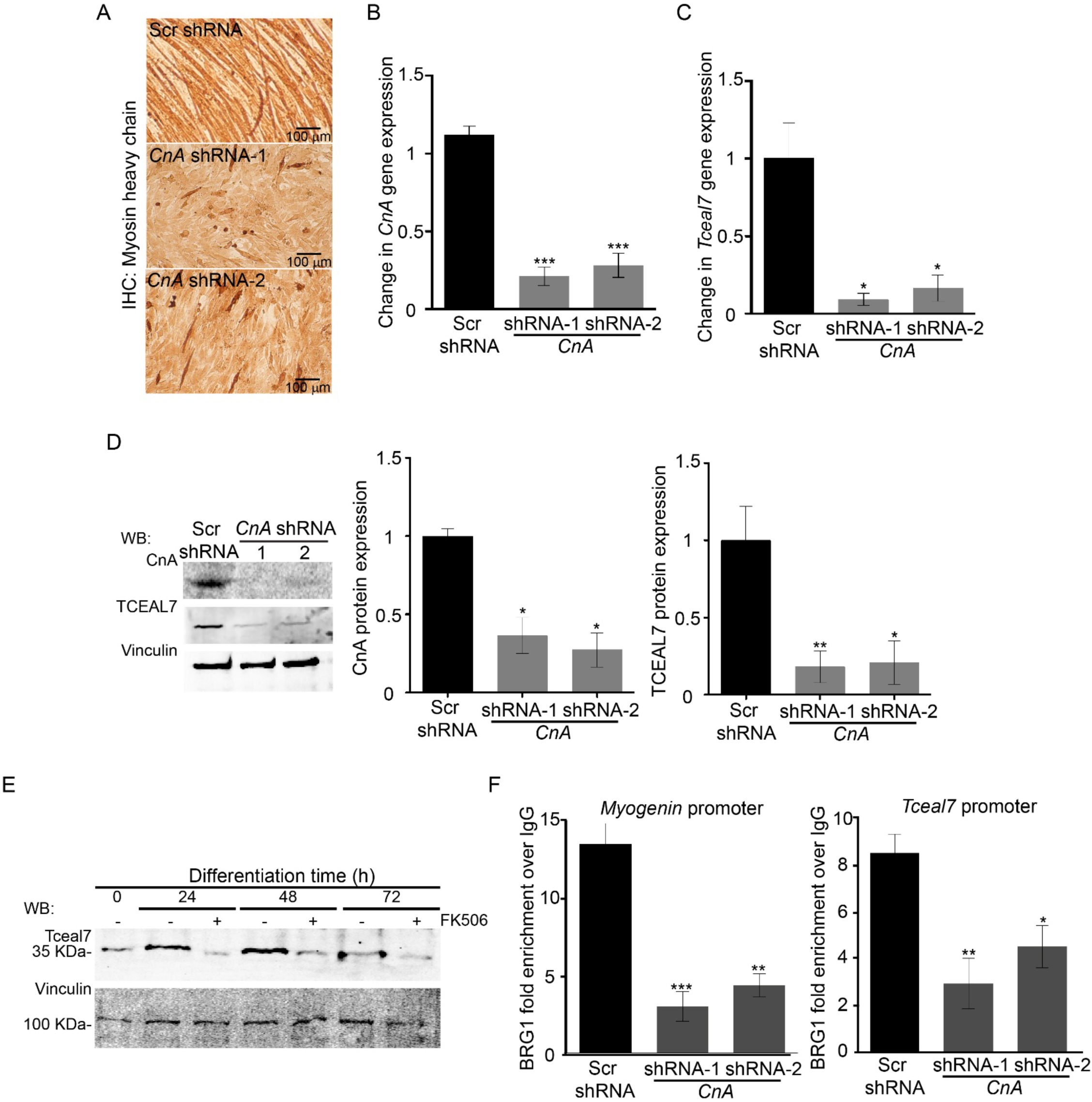
Calcineurin-A knockdown or inhibition impairs myoblast differentiation and suppresses *Tceal7* expression. **(A)** Representative myosin heavy chain immunostaining of 72 h differentiated C2C12 myoblasts transfected with control scrambled sequence (Scr) or one of two distinct siRNAs targeting *CnA*. (**B**) Steady state mRNA expression of *CnA* in 72 h differentiated Scr control and *CnA* knockdown C2C12 myoblasts confirming the efficacy of the *CnA* knockdown. **(C)** Steady state mRNA expression of *Tceal7* at 72 h of differentiation in Scr control and *CnA* knockdown myoblasts. mRNA expression data was normalized to *Eef1a*. **(D)** Representative western blot confirming CnA knockdown and reduced expression of TCEAL7 in 72 h differentiated C2C12 myoblasts. Vinculin was used as the loading control. **(E)** Western blot showing TCEAL7 protein levels in FK506-treated C2C12 cells at the indicated differentiation time points. Vinculin was used as the loading control. **(F)** Chromatin immunoprecipitation showing BRG1 occupancy at the *Myogenin* and *Tceal7* promoters; values were normalized to values obtained for an IgG ChIP. (**F**) Data represents 3 independent biological experiments ± SD. *P < 0.05; ***P < 0.01; ***P < 0.001.

### *Tceal7* is required for myoblast differentiation and regulated myoblast proliferation

Given the differentiation-dependent increase in *Tceal7*, we investigated its functional requirement in myoblast differentiation. shRNA-mediated reduction of *Tceal7* gene expression and protein levels were confirmed by steady state qPCR (**Fig. 4A**) and western blot (**Fig. 4B**). Myoblasts with *Tceal7* knockdown failed to differentiate as demonstrated by reduced MHC staining and the lack of myotubes (**Fig. 4C**).

**Figure 4.**
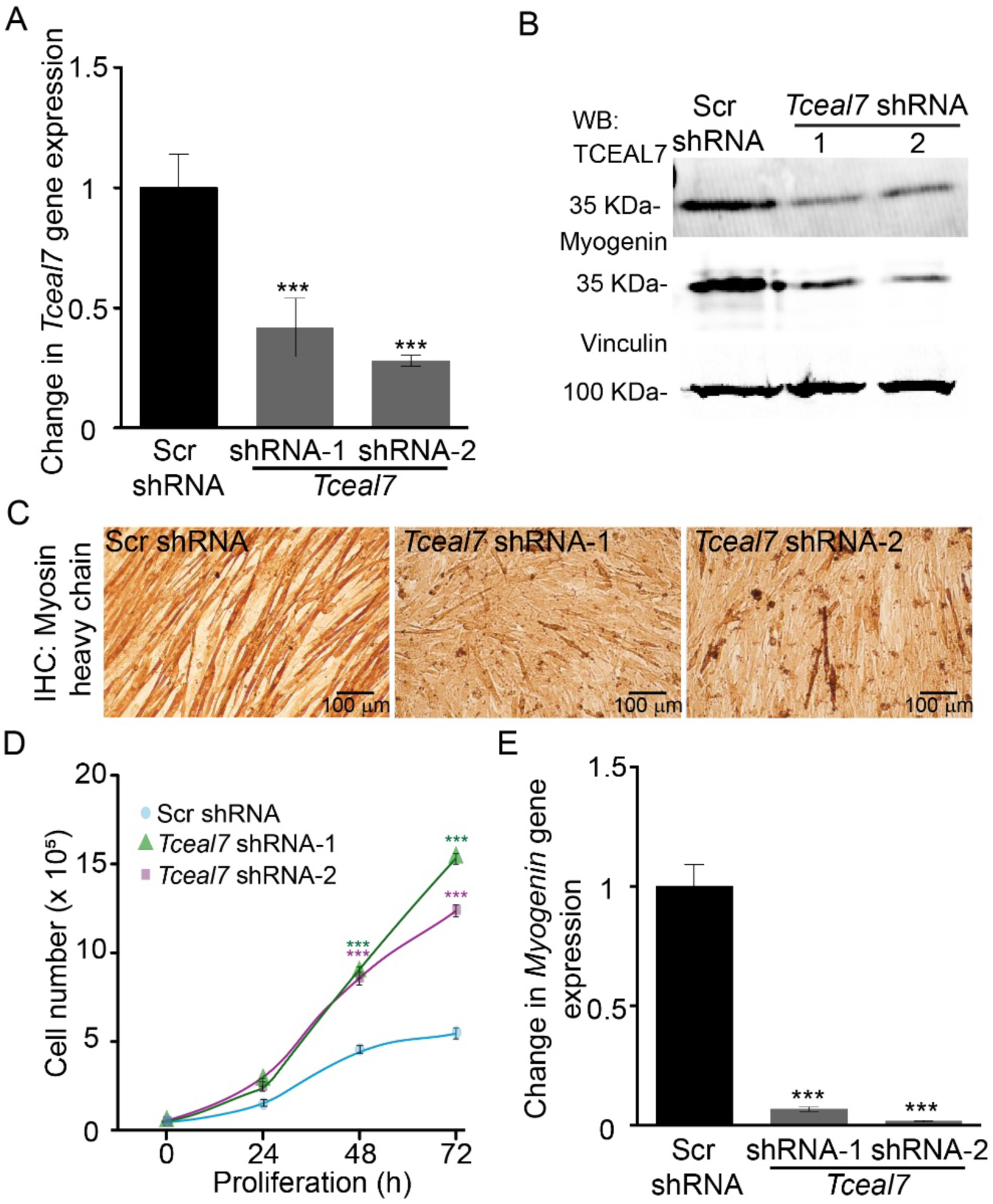
TCEAL7 KD compromises C2C12 myogenic differentiation. **(A)** Steady state mRNA expression levels of *Tceal7* at 72 h of differentiation; the data was normalized to *Eef1a* expression. **(B)** Representative western blot validating siRNA-mediated *Tceal7* knockdown and showing a corresponding decrease in Myogenin expression in 72 h differentiated C2C12 myoblasts. Vinculin was used as loading control. (**C**) Representative light microcopy analyses of myosin heavy chain staining of differentiated C2C12 cells (72 h) transfected with Scr control or *Tceal7* siRNA. (**D**) Cell proliferation analysis comparing *Tceal7* knockdown myoblasts maintained in proliferation conditions with the Scr control myoblasts. **(E)** Steady state mRNA expression levels for *Myogenin* at 72 h differentiation; data were normalized to *Eef1a* expression. Data represents 3 independent biological experiments ± SD. ***P < 0.01.

Prior work demonstrated that *Tceal7* overexpression in C2C12 cells decreased cellular proliferation and increased differentiation capacity [47]. Knockdown of *Tceal7* increased myoblast proliferation (**Fig. 4D**), consistent with prior results. Knockdown of *Tceal7* also reduced Myogenin protein (**Fig. 4B**) and *myogenin* mRNA (**Fig. 4E**) levels, consistent with a requirement for differentiation. TCEAL7 therefore likely contributes to myoblast differentiation by contributing to both cell cycle and to the induction of myogenic gene expression.

### Transcriptomic profiling reveals that *Tceal7* controls differentiation and transcription factor networks

To define the global transcriptional programs regulated by *Tceal7*, we performed RNA-seq on differentiating *Tceal7* knockdown C2C12 myoblasts. Scr control and *Tceal7* knockdown myoblasts were induced to differentiate for 72 h, and total RNA was extracted and processed for RNA-seq analyses. The differentially expressed genes (DEGs) are listed in **Supplemental Table 2**. Gene Ontology (GO) analysis showed that the top ten categories of downregulated genes were all biological processes associated with muscle development, differentiation, and/or function (**Fig. 5A**). Motif enrichment analysis revealed that downregulated genes were highly enriched for motifs associated with several myogenic transcription factors (**Fig. 5B**). The motifs associated with downregulated genes were binding sites for the MRFs, MyoD1, Myf5, and myogenin, and for the E proteins E2A and HEB that heterodimerize with MRFs [53, 72]. These results indicate suppression of the core myogenic regulatory protein network that drives myoblast commitment, differentiation, and fusion. Down regulated genes are also significantly enriched for binding sites for MEF2 family factors (Mef2a, Mef2b, Mef2c, Mef2d), which are cellular transcription factors that augment MRFs to promote transcriptional programs required for sarcomere assembly, metabolic maturation, and myofiber growth [73]. The motifs common to genes that were down regulated by *Tceal7* knockdown reinforce the conclusion that *Tceal7* broadly regulates the myogenic transcription program.

**Figure 5.**
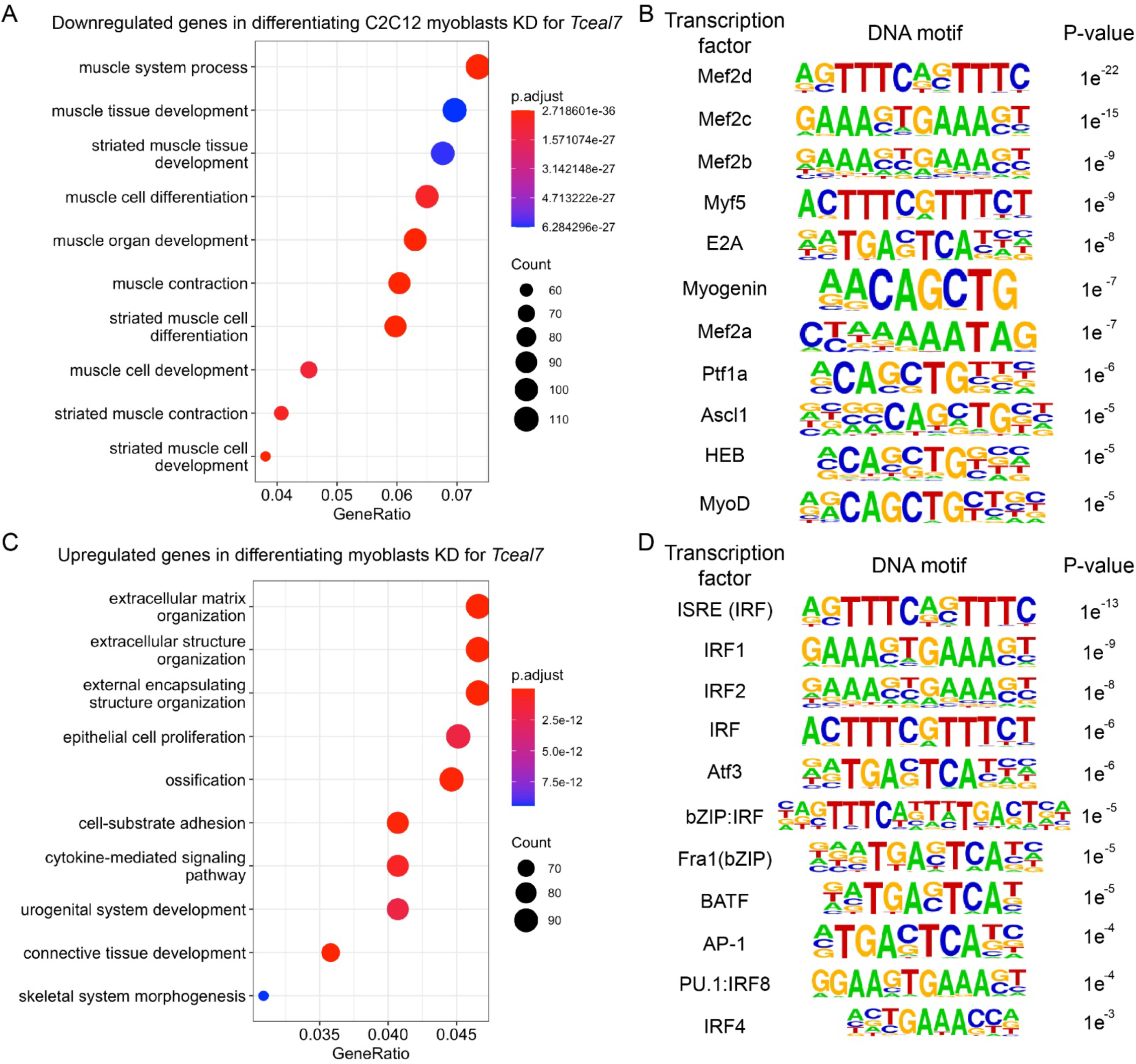
Transcriptomic profiling of differentiated *Tceal7* knockdown C2C12 myoblasts revealed dysregulation of differentiation and cell cycle programs. GO analysis of downregulated (**A**) and upregulated (**C**) genes in differentiated *Tceal7* knockdown myoblasts compared to differentiated Scr-transfected control myoblasts. HOMER known-motif enrichment for downregulated **(B)** and upregulated (**D**) genes.

Upregulated genes were enriched for extracellular matrix structure and organization and cell adhesion. Proliferation and differentiation processes related to other cell types were also identified (**Fig. 5C**). Motif analyses of overexpressed genes (**Fig. 5D**) showed a significant enrichment of ISRE/IRF, IRF1, IRF2, IRF4, and PU.1:IRF8 motifs, suggesting strong activation of interferon-responsive and innate immune transcriptional programs [74]. Identification of motifs for ATF3, BATF, Fra1, and other bZIP/AP-1 factors further point to enhanced stress, cytokine, and inflammatory signaling. Together, the motif profile suggests that reduction of *Tceal7* expression shifts myoblasts toward a heightened immune-stress transcriptional state that is incompatible with effective myogenic differentiation.

PANTHER classification further identified structural and regulatory proteins affected by *Tceal7* knockdown. Among the top categories associated with downregulated genes were pathways that are linked to skeletal muscle development and function (**Fig. 6A**). For instance, the most affected pathway was nicotinic acetylcholine receptor signaling, with 95 genes downregulated. This pathway is essential for neuromuscular junction formation and activity-dependent maturation of myofibers, promoting terminal differentiation and sarcomere organization [75]. Seventy-seven genes related to cytoskeletal regulation by Rho GTPases, which controls actin dynamics, myoblast shape, migration, and the membrane remodeling required for myoblast alignment and fusion were repressed by *Tceal7* knockdown. Chemokine- and cytokine-mediated inflammation pathways showed decreased expression of 259 related genes. These inflammatory pathways are known to regulate the inflammatory milieu required for proper muscle regeneration, balancing early pro-inflammatory cues that promote myoblast activation with later anti-inflammatory signals that enable differentiation and fusion [76]. β_1-_ and β_2_-adrenergic receptor signaling, which broadly impacts muscle and muscle stem cell function [77–79] were also significantly affected with 45 genes downregulated. The PI3K pathway is a central driver of myogenesis, integrating growth factor cues to stimulate Akt signaling, protein synthesis, survival, and myotube growth [80]. Fifty-three genes associated with PI3K were repressed. Finally, the Wnt signaling pathway, with 307 genes downregulated, represented the largest number of genes altered. The Wnt pathway is a well-established regulator of skeletal myogenesis, directing progenitor cell fate, initiating differentiation programs, and coordinating myotube formation and growth [41, 42, 81].

**Figure 6:**
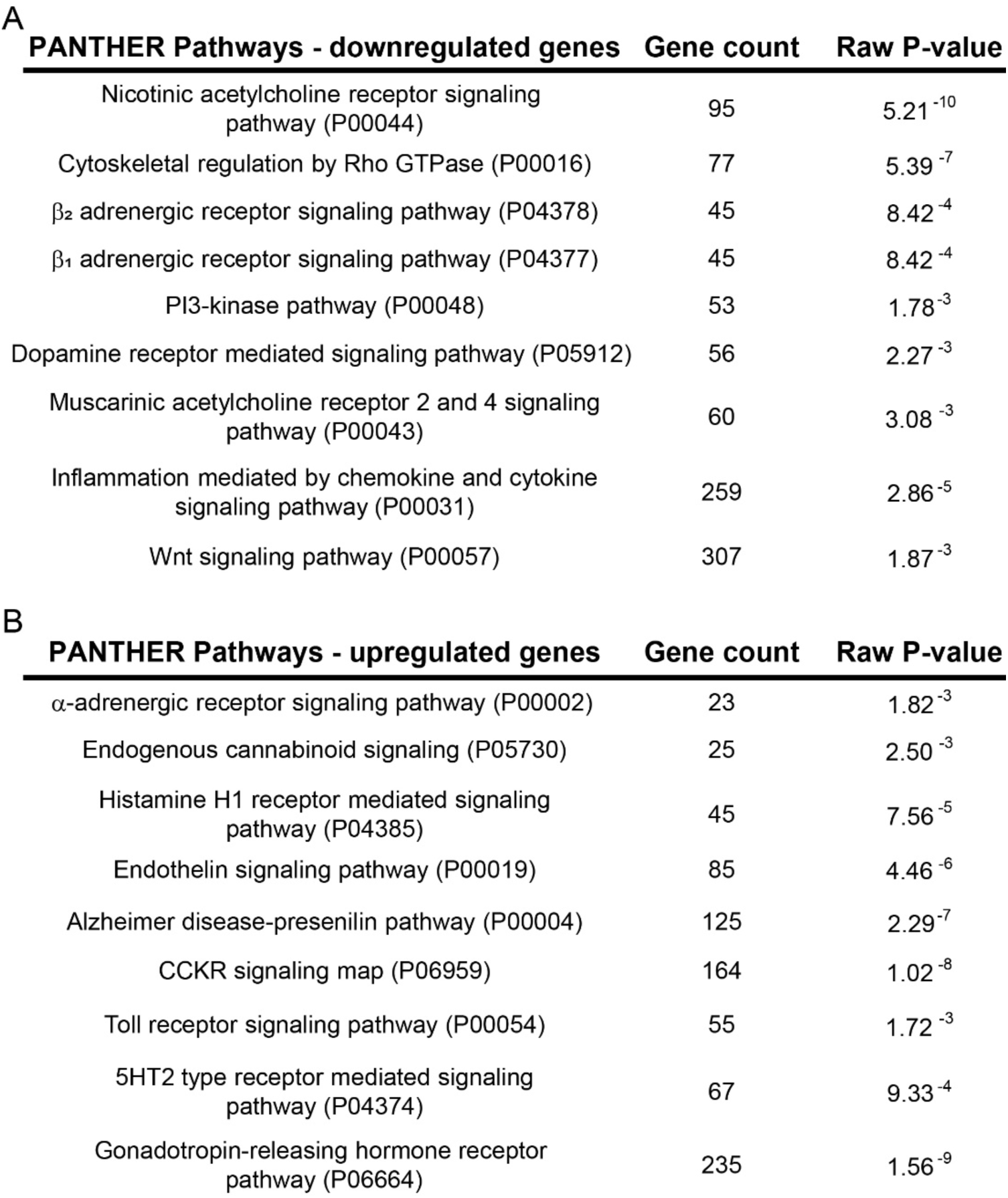
PANTHER analysis top terms for down regulated and up regulated genes in differentiated *Tceal7* knockdown myoblasts.

PANTHER analyses also provided insight into upregulated pathways, identifying signaling programs that may impact skeletal muscle development, regeneration or function either directly or indirectly (**Fig. 6B**). Among these was endothelin receptor signaling, recently shown to inhibit myoblast proliferation and MRF and other myogenic gene expression through the p38 MAPK pathway. Chronic exposure to endothelin led to skeletal muscle atrophy and poor exercise performance [82], Endothelin also promotes muscle fibrosis [83]. α-adrenergic receptor, a G-protein coupled receptor best known for its role in vasoconstriction, indirectly affects muscle function [84] but may have direct effects as well, as chemical mimics of pathway components resulted in depletion of a histone deacetylase from the nucleus of slow-twitch soleus muscle fibers, resulting in altered gene regulation [85]. Endogenous cannabinoid signaling uses lipid messenger molecules to regulate numerous body functions. Levels of an endogenous cannabinoid normally decrease during satellite cell and myoblast differentiation, and its continued presence inhibits differentiation [86]. Thus, upregulation of this pathway by *Tceal7* knockdown correlates well with the observed inhibition of myoblast differentiation observed upon *Tceal7* knockdown. Finally, presenilin knockdown led to accelerated myoblast differentiation during muscle regeneration in vivo, while constitutive presenilin expression suppressed differentiation [87].

Collectively, the PANTHER analysis confirms the widespread impact of *Tceal7* knockdown on the fidelity of gene expression in myoblasts, with broad effects across a spectrum of genes and signaling pathways.

## DISCUSSION

Collectively, these results identify *Tceal7* as a BRG1-dependent, calcineurin-responsive regulator of myogenic differentiation. *Tceal7* is induced during differentiation, localizes to both the nucleus and the cytoplasm, and is required for myogenin expression, myotube fusion, and activation of myogenic gene networks. *Tceal7* knockdown also upregulates GPCR-, cytokine-, and interferon-related pathways and IRF/AP-1 responsive-genes, shifting cells toward an inflammatory stress state that are likely incompatible with differentiation. Our results support a model in which calcineurin signaling promotes BRG1 recruitment to the *Tceal7* promoter, enabling its expression and activation of the downstream transcriptional environment required for efficient myogenic commitment and skeletal muscle formation, maturation, and function.

The TCEAL gene family comprises a cluster of X-linked genes conserved across mammals. Members of this family were initially presumed to function as transcription elongation factors based on early reports that TCEAL1 shared limited homology with TFIIS, an RNA polymerase II elongation factor containing a characteristic Zn-binding domain [3]. However, structural comparisons using AlphaFold2 have since demonstrated that TCEAL proteins lack the Zn-binding and Pol II-interacting domains of TFIIS, and no biochemical evidence supports their direct participation in transcriptional elongation or RNA polymerase II binding [10]. Thus, the molecular activities of TCEAL proteins remain largely undefined. It remains possible that TCEAL7 plays a direct role in regulating transcription. It is also possible that it indirectly affects gene expression, perhaps through the de-ubiquitination function identified in other family members [11–13] or in an as yet undiscovered function. A more general and indirect mechanism of action may make sense, given the extensive effect seen in the analyses of the RNA-seq data from the *Tceal7* knockdown myoblasts where genes associated with a multitude of muscle and muscle-related functions and pathways were mis-regulated. The wide-spread association between mis-regulation of genes encoding TCEAL family members with a large number of disparate types of cancers [10] may also argue that TCEAL7 and other family members have a general and indirect impact on gene expression.

The regulation of the *Tceal7* gene during embryonic development and during in vitro myoblast differentiation established that MyoD and related MRFs, along with cooperating factors, regulate transcription in muscle [46–50]. Our work extends these findings to demonstrate that the mSWI/SNF chromatin remodeling enzymes containing BRG1 as the ATPase subunit also regulate the expression of *Tceal7*. We have previously shown that BRG1 remodels chromatin at target genes to facilitate stable binding of MRFs [28]; it is likely that this function extends to the *Tceal7* promoter. In addition, we demonstrated that BRG1 function is sensitive to chemical inhibition of reduction in expression of the Cn phosphatase and demonstrated that dephosphorylation of BRG1 was associated with its activation at target promoters [69]. In keeping with the prior model, we propose that Cn-dependent modification of BRG1 promotes stable binding of MRFs at the *Tceal7* promoter, thereby facilitating gene induction during myoblast differentiation. It remains to be seen whether regulation of *Tceal7* gene expression by Cn and BRG1 regulation occurs in other cell types. Another interesting question is whether Cn and BRG1 mediate regulation of the genes encoding other TCEAL family members.

## CONCLUSIONS

In summary, this study identifies TCEAL7 as a critical downstream effector of BRG1-dependent chromatin remodeling and calcineurin signaling during myoblast differentiation. We propose a model in which *Tceal7* expression is facilitated by BRG1 recruitment to the *Tceal7* promoter, where its chromatin remodeling activity is activated by calcineurin phosphatase, resulting in the stable association of MRFs and other cooperating transcription factors that drive myogenic gene expression. We also conclude that TCEAL7 plays an essential role in coordinating signaling pathways, cell cycle exit, the induction of the myogenic transcription programs, and efficient myoblast fusion into myotubes. Moreover, our work demonstrates that the spectrum of TCEAL7 function is extremely broad, impacting nearly every aspect of myoblast differentiation. Additionally, we suggest that TCEAL7 function will be generally required for not only for tissue differentiation and development but also for cell proliferation and for cell and tissue homestasis.

## FUNDING

Research reported in this publication was supported by the National Institutes of Health under Award Number R35GM136393 to ANI and R01AR077578 to TPB. The content is solely the responsibility of the authors and does not necessarily represent the official views of the National Institutes of Health.

## AUTHOR CONTRIBUTIONS

SY, TS, and ANI conceived the project. SY generated all of the data. TS performed the bioinformatic analyses. All four authors analyzed data. SY, TS, and TP-B prepared tables and figures. TP-B and ANI drafted the manuscript. All four authors reviewed and revised the manuscript.

## Notes

### Competing Interest Statement

The authors have declared no competing interest.

## REFERENCES

1. Navas-Perez E, Vicente-Garcia C, Mirra S, Burguera D, Fernandez-Castillo N, Ferran JL, et al. Characterization of an eutherian gene cluster generated after transposon domestication identifies Bex3 as relevant for advanced neurological functions. Genome Biol. 2020;21(1):267. Epub 2020/10/27. doi: 10.1186/s13059-020-02172-3. PubMed PMID: 33100228; PubMed Central PMCID: PMCPMC7586669.

2. Winter EE, Ponting CP. Mammalian BEX, WEX and GASP genes: coding and non-coding chimaerism sustained by gene conversion events. BMC Evol Biol. 2005;5:54. Epub 2005/10/14. doi: 10.1186/1471-2148-5-54. PubMed PMID: 16221301; PubMed Central PMCID: PMCPMC1274310.

3. Yeh CH, Shatkin AJ. A HeLa-cell-encoded p21 is homologous to transcription elongation factor SII. Gene. 1994;143(2):285–7. Epub 1994/06/10. doi: 10.1016/0378-1119(94)90112-0. PubMed PMID: 8206389.

4. Natori S, Takeuchi K, Takahashi K, Mizuno D. DNA dependent RNA polymerase from Ehrlich ascites tumor cells. II. Factors stimulating the activity of RNA polymerase II. J Biochem. 1973;73(4):879–88. Epub 1973/04/01. doi: 10.1093/oxfordjournals.jbchem.a130150. PubMed PMID: 4352629.

5. Reines D, Conaway JW, Conaway RC. The RNA polymerase II general elongation factors. Trends Biochem Sci. 1996;21(9):351–5. Epub 1996/09/01. PubMed PMID: 8870500; PubMed Central PMCID: PMCPMC3374595.

6. Sekimizu K, Kobayashi N, Mizuno D, Natori S. Purification of a factor from Ehrlich ascites tumor cells specifically stimulating RNA polymerase II. Biochemistry. 1976;15(23):5064–70. Epub 1976/11/16. doi: 10.1021/bi00668a018. PubMed PMID: 990265.

7. Wind M, Reines D. Transcription elongation factor SII. Bioessays. 2000;22(4):327–36. Epub 2000/03/21. doi: 10.1002/(SICI)1521-1878(200004)22:4<327::AID-BIES3>3.0.CO;2-4. PubMed PMID: 10723030; PubMed Central PMCID: PMCPMC3367499.

8. Jumper J, Evans R, Pritzel A, Green T, Figurnov M, Ronneberger O, et al. Highly accurate protein structure prediction with AlphaFold. Nature. 2021;596(7873):583–9. Epub 2021/07/16. doi: 10.1038/s41586-021-03819-2. PubMed PMID: 34265844; PubMed Central PMCID: PMCPMC8371605 have filed non-provisional patent applications 16/701,070 and PCT/EP2020/084238, and provisional patent applications 63/107,362, 63/118,917, 63/118,918, 63/118,921 and 63/118,919, each in the name of DeepMind Technologies Limited, each pending, relating to machine learning for predicting protein structures. The other authors declare no competing interests.

9. Yang Z, Zeng X, Zhao Y, Chen R. AlphaFold2 and its applications in the fields of biology and medicine. Signal Transduct Target Ther. 2023;8(1):115. Epub 2023/03/16. doi: 10.1038/s41392-023-01381-z. PubMed PMID: 36918529; PubMed Central PMCID: PMCPMC10011802.

10. Yadav S, Imbalzano AN. Transcription Elongation Factor A (SII)-Like (TCEAL) Family Members: Re-Evaluation of Predicted Structure and Summary of Expression and Functions. Crit Rev Eukaryot Gene Expr. 2025;35(8):11–33. doi: 10.1615/CritRevEukaryotGeneExpr.2025061337. PubMed PMID: 41427930.

11. Hermanns T, Pichlo C, Woiwode I, Klopffleisch K, Witting KF, Ovaa H, et al. A family of unconventional deubiquitinases with modular chain specificity determinants. Nat Commun. 2018;9(1):799. Epub 2018/02/25. doi: 10.1038/s41467-018-03148-5. PubMed PMID: 29476094; PubMed Central PMCID: PMCPMC5824887.

12. Querques F, Darling S, Cheetham-Wilkinson I, Kim RQ, Hapangama DK, Sixma TK, et al. S18-phosphorylation of USP7 regulates interaction with TCEAL4 that defines specific complexes and potentially distinct functions. bioRxiv. 2021:2021.07.07.451439. doi: 10.1101/2021.07.07.451439.

13. Szklarczyk D, Kirsch R, Koutrouli M, Nastou K, Mehryary F, Hachilif R, et al. The STRING database in 2023: protein-protein association networks and functional enrichment analyses for any sequenced genome of interest. Nucleic Acids Res. 2023;51(D1):D638–D46. Epub 2022/11/13. doi: 10.1093/nar/gkac1000. PubMed PMID: 36370105; PubMed Central PMCID: PMCPMC9825434.

14. Imbalzano AN, Kwon H, Green MR, Kingston RE. Facilitated binding of TATA-binding protein to nucleosomal DNA. Nature. 1994;370(6489):481–5. doi: 10.1038/370481a0. PubMed PMID: 8047170.

15. Kwon H, Imbalzano AN, Khavari PA, Kingston RE, Green MR. Nucleosome disruption and enhancement of activator binding by a human SW1/SNF complex. Nature. 1994;370(6489):477–81. doi: 10.1038/370477a0. PubMed PMID: 8047169.

16. Wang W, Cote J, Xue Y, Zhou S, Khavari PA, Biggar SR, et al. Purification and biochemical heterogeneity of the mammalian SWI-SNF complex. EMBO J. 1996;15(19):5370–82. PubMed PMID: 8895581; PubMed Central PMCID: PMCPMC452280.

17. Hargreaves DC, Crabtree GR. ATP-dependent chromatin remodeling: genetics, genomics and mechanisms. Cell Res. 2011;21(3):396–420. Epub 20110301. doi: 10.1038/cr.2011.32. PubMed PMID: 21358755; PubMed Central PMCID: PMCPMC3110148.

18. Wu JI. Diverse functions of ATP-dependent chromatin remodeling complexes in development and cancer. Acta Biochim Biophys Sin (Shanghai). 2012;44(1):54–69. doi: 10.1093/abbs/gmr099. PubMed PMID: 22194014.

19. Mashtalir N, D’Avino AR, Michel BC, Luo J, Pan J, Otto JE, et al. Modular Organization and Assembly of SWI/SNF Family Chromatin Remodeling Complexes. Cell. 2018;175(5):1272–88 e20. Epub 20181018. doi: 10.1016/j.cell.2018.09.032. PubMed PMID: 30343899; PubMed Central PMCID: PMCPMC6791824.

20. Barutcu AR, Lajoie BR, Fritz AJ, McCord RP, Nickerson JA, van Wijnen AJ, et al. SMARCA4 regulates gene expression and higher-order chromatin structure in proliferating mammary epithelial cells. Genome Res. 2016;26(9):1188–201. Epub 20160719. doi: 10.1101/gr.201624.115. PubMed PMID: 27435934; PubMed Central PMCID: PMCPMC5052043.

21. Brownlee PM, Meisenberg C, Downs JA. The SWI/SNF chromatin remodelling complex: Its role in maintaining genome stability and preventing tumourigenesis. DNA Repair (Amst). 2015;32:127–33. Epub 20150501. doi: 10.1016/j.dnarep.2015.04.023. PubMed PMID: 25981841.

22. Ribeiro-Silva C, Vermeulen W, Lans H. SWI/SNF: Complex complexes in genome stability and cancer. DNA Repair (Amst). 2019;77:87–95. Epub 20190315. doi: 10.1016/j.dnarep.2019.03.007. PubMed PMID: 30897376.

23. Takebayashi S, Lei I, Ryba T, Sasaki T, Dileep V, Battaglia D, et al. Murine esBAF chromatin remodeling complex subunits BAF250a and Brg1 are necessary to maintain and reprogram pluripotency-specific replication timing of select replication domains. Epigenetics Chromatin. 2013;6(1):42. Epub 20131213. doi: 10.1186/1756-8935-6-42. PubMed PMID: 24330833; PubMed Central PMCID: PMCPMC3895691.

24. Kadoch C, Crabtree GR. Mammalian SWI/SNF chromatin remodeling complexes and cancer: Mechanistic insights gained from human genomics. Sci Adv. 2015;1(5):e1500447. Epub 20150612. doi: 10.1126/sciadv.1500447. PubMed PMID: 26601204; PubMed Central PMCID: PMCPMC4640607.

25. Kadoch C, Hargreaves DC, Hodges C, Elias L, Ho L, Ranish J, et al. Proteomic and bioinformatic analysis of mammalian SWI/SNF complexes identifies extensive roles in human malignancy. Nat Genet. 2013;45(6):592–601. Epub 20130505. doi: 10.1038/ng.2628. PubMed PMID: 23644491; PubMed Central PMCID: PMCPMC3667980.

26. Shain AH, Pollack JR. The spectrum of SWI/SNF mutations, ubiquitous in human cancers. PLoS One. 2013;8(1):e55119. Epub 20130123. doi: 10.1371/journal.pone.0055119. PubMed PMID: 23355908; PubMed Central PMCID: PMCPMC3552954.

27. de la Serna IL, Carlson KA, Imbalzano AN. Mammalian SWI/SNF complexes promote MyoD-mediated muscle differentiation. Nat Genet. 2001;27(2):187–90. doi: 10.1038/84826. PubMed PMID: 11175787.

28. de la Serna IL, Ohkawa Y, Berkes CA, Bergstrom DA, Dacwag CS, Tapscott SJ, et al. MyoD targets chromatin remodeling complexes to the myogenin locus prior to forming a stable DNA-bound complex. Mol Cell Biol. 2005;25(10):3997–4009. doi: 10.1128/MCB.25.10.3997-4009.2005. PubMed PMID: 15870273; PubMed Central PMCID: PMCPMC1087700.

29. Han S, Jin L, Peng W, Lv X, Zhang Z, Liu T, et al. Baf60c in skeletal muscle regulates adipose tissue thermogenesis via Musclin-mediated endocrine signaling. Life Metab. 2025;4(4):loaf015. Epub 20250504. doi: 10.1093/lifemeta/loaf015. PubMed PMID: 40585527; PubMed Central PMCID: PMCPMC12201986.

30. Kang JS, Kim D, Rhee J, Seo JY, Park I, Kim JH, et al. Baf155 regulates skeletal muscle metabolism via HIF-1a signaling. PLoS Biol. 2023;21(7):e3002192. Epub 20230721. doi: 10.1371/journal.pbio.3002192. PubMed PMID: 37478146; PubMed Central PMCID: PMCPMC10396025.

31. Meng ZX, Li S, Wang L, Ko HJ, Lee Y, Jung DY, et al. Baf60c drives glycolytic metabolism in the muscle and improves systemic glucose homeostasis through Deptor-mediated Akt activation. Nat Med. 2013;19(5):640–5. Epub 20130407. doi: 10.1038/nm.3144. PubMed PMID: 23563706; PubMed Central PMCID: PMCPMC3650110.

32. Sharma T, Robinson DCL, Witwicka H, Dilworth FJ, Imbalzano AN. The Bromodomains of the mammalian SWI/SNF (mSWI/SNF) ATPases Brahma (BRM) and Brahma Related Gene 1 (BRG1) promote chromatin interaction and are critical for skeletal muscle differentiation. Nucleic Acids Res. 2021;49(14):8060–77. doi: 10.1093/nar/gkab617. PubMed PMID: 34289068; PubMed Central PMCID: PMCPMC8373147.

33. Ohkawa Y, Yoshimura S, Higashi C, Marfella CG, Dacwag CS, Tachibana T, et al. Myogenin and the SWI/SNF ATPase Brg1 maintain myogenic gene expression at different stages of skeletal myogenesis. J Biol Chem. 2007;282(9):6564–70. Epub 20061227. doi: 10.1074/jbc.M608898200. PubMed PMID: 17194702.

34. Khavari PA, Peterson CL, Tamkun JW, Mendel DB, Crabtree GR. BRG1 contains a conserved domain of the SWI2/SNF2 family necessary for normal mitotic growth and transcription. Nature. 1993;366(6451):170–4. doi: 10.1038/366170a0. PubMed PMID: 8232556.

35. Reyes JC, Barra J, Muchardt C, Camus A, Babinet C, Yaniv M. Altered control of cellular proliferation in the absence of mammalian brahma (SNF2alpha). EMBO J. 1998;17(23):6979–91. doi: 10.1093/emboj/17.23.6979. PubMed PMID: 9843504; PubMed Central PMCID: PMCPMC1171046.

36. Kowenz-Leutz E, Leutz A. A C/EBP beta isoform recruits the SWI/SNF complex to activate myeloid genes. Mol Cell. 1999;4(5):735–43. doi: 10.1016/s1097-2765(00)80384-6. PubMed PMID: 10619021.

37. Pedersen TA, Kowenz-Leutz E, Leutz A, Nerlov C. Cooperation between C/EBPalpha TBP/TFIIB and SWI/SNF recruiting domains is required for adipocyte differentiation. Genes Dev. 2001;15(23):3208–16. doi: 10.1101/gad.209901. PubMed PMID: 11731483; PubMed Central PMCID: PMCPMC312836.

38. Barker N, Hurlstone A, Musisi H, Miles A, Bienz M, Clevers H. The chromatin remodelling factor Brg-1 interacts with beta-catenin to promote target gene activation. EMBO J. 2001;20(17):4935–43. doi: 10.1093/emboj/20.17.4935. PubMed PMID: 11532957; PubMed Central PMCID: PMCPMC125268.

39. Griffin CT, Curtis CD, Davis RB, Muthukumar V, Magnuson T. The chromatin-remodeling enzyme BRG1 modulates vascular Wnt signaling at two levels. Proc Natl Acad Sci U S A. 2011;108(6):2282–7. Epub 20110124. doi: 10.1073/pnas.1013751108. PubMed PMID: 21262838; PubMed Central PMCID: PMCPMC3038709.

40. Wagner G, Singhal N, Nicetto D, Straub T, Kremmer E, Rupp RAW. Brg1 chromatin remodeling ATPase balances germ layer patterning by amplifying the transcriptional burst at midblastula transition. PLoS Genet. 2017;13(5):e1006757. Epub 20170512. doi: 10.1371/journal.pgen.1006757. PubMed PMID: 28498870; PubMed Central PMCID: PMCPMC5428918.

41. Munsterberg AE, Kitajewski J, Bumcrot DA, McMahon AP, Lassar AB. Combinatorial signaling by Sonic hedgehog and Wnt family members induces myogenic bHLH gene expression in the somite. Genes Dev. 1995;9(23):2911–22. doi: 10.1101/gad.9.23.2911. PubMed PMID: 7498788.

42. Stern HM, Brown AM, Hauschka SD. Myogenesis in paraxial mesoderm: preferential induction by dorsal neural tube and by cells expressing Wnt-1. Development. 1995;121(11):3675–86. doi: 10.1242/dev.121.11.3675. PubMed PMID: 8582280.

43. Fedorov O, Castex J, Tallant C, Owen DR, Martin S, Aldeghi M, et al. Selective targeting of the BRG/PB1 bromodomains impairs embryonic and trophoblast stem cell maintenance. Sci Adv. 2015;1(10):e1500723. Epub 20151113. doi: 10.1126/sciadv.1500723. PubMed PMID: 26702435; PubMed Central PMCID: PMCPMC4681344.

44. Gerstenberger BS, Trzupek JD, Tallant C, Fedorov O, Filippakopoulos P, Brennan PE, et al. Identification of a Chemical Probe for Family VIII Bromodomains through Optimization of a Fragment Hit. J Med Chem. 2016;59(10):4800–11. Epub 20160503. doi: 10.1021/acs.jmedchem.6b00012. PubMed PMID: 27115555; PubMed Central PMCID: PMCPMC5034155.

45. Sharma T, Olea-Flores M, Imbalzano AN. Regulation of the Wnt signaling pathway during myogenesis by the mammalian SWI/SNF ATPase BRG1. Front Cell Dev Biol. 2023;11:1160227. Epub 20230707. doi: 10.3389/fcell.2023.1160227. PubMed PMID: 37484913; PubMed Central PMCID: PMCPMC10360407.

46. Sawada A, Yamamoto T, Sato T. Tceal5 and Tceal7 Function in C2C12 Myogenic Differentiation via Exosomes in Fetal Bovine Serum. Int J Mol Sci. 2022;23(4). Epub 2022/02/27. doi: 10.3390/ijms23042036. PubMed PMID: 35216152; PubMed Central PMCID: PMCPMC8877866.

47. Shi X, Garry DJ. Myogenic regulatory factors transactivate the Tceal7 gene and modulate muscle differentiation. Biochem J. 2010;428(2):213–21. Epub 2010/03/24. doi: 10.1042/BJ20091906. PubMed PMID: 20307260.

48. Xiong Z, Wang M, Wu J, Shi X. Tceal7 Regulates Skeletal Muscle Development through Its Interaction with Cdk1. Int J Mol Sci. 2023;24(7). Epub 2023/04/14. doi: 10.3390/ijms24076264. PubMed PMID: 37047236; PubMed Central PMCID: PMCPMC10094454.

49. Xiong Z, Wang M, You S, Chen X, Lin J, Wu J, et al. Transcription Regulation of Tceal7 by the Triple Complex of Mef2c, Creb1 and Myod. Biology (Basel). 2022;11(3). Epub 2022/03/27. doi: 10.3390/biology11030446. PubMed PMID: 35336819; PubMed Central PMCID: PMCPMC8945367.

50. Rana K, Lee NK, Zajac JD, MacLean HE. Expression of androgen receptor target genes in skeletal muscle. Asian J Androl. 2014;16(5):675–83. doi: 10.4103/1008-682X.122861. PubMed PMID: 24713826; PubMed Central PMCID: PMCPMC4215656.

51. Hulmi JJ, Nissinen TA, Rasanen M, Degerman J, Lautaoja JH, Hemanthakumar KA, et al. Prevention of chemotherapy-induced cachexia by ACVR2B ligand blocking has different effects on heart and skeletal muscle. J Cachexia Sarcopenia Muscle. 2018;9(2):417–32. Epub 20171211. doi: 10.1002/jcsm.12265. PubMed PMID: 29230965; PubMed Central PMCID: PMCPMC5879968.

52. Hong AR, Kim K, Lee JY, Yang JY, Kim JH, Shin CS, et al. Transformation of Mature Osteoblasts into Bone Lining Cells and RNA Sequencing-Based Transcriptome Profiling of Mouse Bone during Mechanical Unloading. Endocrinol Metab (Seoul). 2020;35(2):456–69. Epub 20200624. doi: 10.3803/EnM.2020.35.2.456. PubMed PMID: 32615730; PubMed Central PMCID: PMCPMC7386115.

53. Zammit PS. Function of the myogenic regulatory factors Myf5, MyoD, Myogenin and MRF4 in skeletal muscle, satellite cells and regenerative myogenesis. Semin Cell Dev Biol. 2017;72:19–32. Epub 20171115. doi: 10.1016/j.semcdb.2017.11.011. PubMed PMID: 29127046.

54. Polyak K, Lee MH, Erdjument-Bromage H, Koff A, Roberts JM, Tempst P, et al. Cloning of p27Kip1, a cyclin-dependent kinase inhibitor and a potential mediator of extracellular antimitogenic signals. Cell. 1994;78(1):59–66. doi: 10.1016/0092-8674(94)90572-x. PubMed PMID: 8033212.

55. Gupta J, Saeed BI, Bishoyi AK, Alkhathami AG, Asliddin S, Nathiya D, et al. From cell cycle control to cancer therapy: exploring the role of CDK1 and CDK2 in tumorigenesis. Med Oncol. 2025;42(9):422. Epub 20250809. doi: 10.1007/s12032-025-02973-1. PubMed PMID: 40782258.

56. Blau HM, Chiu CP, Webster C. Cytoplasmic activation of human nuclear genes in stable heterocaryons. Cell. 1983;32(4):1171–80. doi: 10.1016/0092-8674(83)90300-8. PubMed PMID: 6839359.

57. Witwicka H, Nogami J, Syed SA, Maehara K, Padilla-Benavides T, Ohkawa Y, et al. Calcineurin Broadly Regulates the Initiation of Skeletal Muscle-Specific Gene Expression by Binding Target Promoters and Facilitating the Interaction of the SWI/SNF Chromatin Remodeling Enzyme. Mol Cell Biol. 2019;39(19). Epub 20190911. doi: 10.1128/MCB.00063-19. PubMed PMID: 31308130; PubMed Central PMCID: PMCPMC6751634.

58. Schneider CA, Rasband WS, Eliceiri KW. NIH Image to ImageJ: 25 years of image analysis. Nat Methods. 2012;9(7):671–5. doi: 10.1038/nmeth.2089. PubMed PMID: 22930834; PubMed Central PMCID: PMCPMC5554542.

59. Livak KJ, Schmittgen TD. Analysis of relative gene expression data using real-time quantitative PCR and the 2(-Delta Delta C(T)) Method. Methods. 2001;25(4):402–8. doi: 10.1006/meth.2001.1262. PubMed PMID: 11846609.

60. Liao Y, Smyth GK, Shi W. featureCounts: an efficient general purpose program for assigning sequence reads to genomic features. Bioinformatics. 2014;30(7):923–30. Epub 20131113. doi: 10.1093/bioinformatics/btt656. PubMed PMID: 24227677.

61. Love MI, Huber W, Anders S. Moderated estimation of fold change and dispersion for RNA-seq data with DESeq2. Genome Biol. 2014;15(12):550. doi: 10.1186/s13059-014-0550-8. PubMed PMID: 25516281; PubMed Central PMCID: PMCPMC4302049.

62. Wu T, Hu E, Xu S, Chen M, Guo P, Dai Z, et al. clusterProfiler 4.0: A universal enrichment tool for interpreting omics data. Innovation (Camb). 2021;2(3):100141. Epub 20210701. doi: 10.1016/j.xinn.2021.100141. PubMed PMID: 34557778; PubMed Central PMCID: PMCPMC8454663.

63. Mi H, Vandergriff J, Campbell M, Narechania A, Majoros W, Lewis S, et al. Assessment of genome-wide protein function classification for Drosophila melanogaster. Genome Res. 2003;13(9):2118–28. doi: 10.1101/gr.771603. PubMed PMID: 12952880; PubMed Central PMCID: PMCPMC403707.

64. Thomas PD, Kejariwal A, Campbell MJ, Mi H, Diemer K, Guo N, et al. PANTHER: a browsable database of gene products organized by biological function, using curated protein family and subfamily classification. Nucleic Acids Res. 2003;31(1):334–41. doi: 10.1093/nar/gkg115. PubMed PMID: 12520017; PubMed Central PMCID: PMCPMC165562.

65. Guo Y, Liao Y, Jia C, Ren J, Wang J, Li T. MicroRNA-182 promotes tumor cell growth by targeting transcription elongation factor A-like 7 in endometrial carcinoma. Cell Physiol Biochem. 2013;32(3):581–90. Epub 2013/09/12. doi: 10.1159/000354462. PubMed PMID: 24021963.

66. Yue X, Lan F, Xia T. Hypoxic Glioma Cell-Secreted Exosomal miR-301a Activates Wnt/beta-catenin Signaling and Promotes Radiation Resistance by Targeting TCEAL7. Mol Ther. 2019;27(11):1939–49. Epub 2019/08/14. doi: 10.1016/j.ymthe.2019.07.011. PubMed PMID: 31402274; PubMed Central PMCID: PMCPMC6838947.

67. Huang CY, Chen YM, Zhao JJ, Chen YB, Jiang SS, Yan SM, et al. Decreased expression of transcription elongation factor A-like 7 is associated with gastric adenocarcinoma prognosis. PLoS One. 2013;8(1):e54671. Epub 2013/02/02. doi: 10.1371/journal.pone.0054671. PubMed PMID: 23372750; PubMed Central PMCID: PMCPMC3555988.

68. Simone C, Forcales SV, Hill DA, Imbalzano AN, Latella L, Puri PL. p38 pathway targets SWI-SNF chromatin-remodeling complex to muscle-specific loci. Nat Genet. 2004;36(7):738–43. Epub 20040620. doi: 10.1038/ng1378. PubMed PMID: 15208625.

69. Nasipak BT, Padilla-Benavides T, Green KM, Leszyk JD, Mao W, Konda S, et al. Opposing calcium-dependent signalling pathways control skeletal muscle differentiation by regulating a chromatin remodelling enzyme. Nat Commun. 2015;6:7441. Epub 20150617. doi: 10.1038/ncomms8441. PubMed PMID: 26081415; PubMed Central PMCID: PMCPMC4530624.

70. Klee CB, Crouch TH, Krinks MH. Calcineurin: a calcium- and calmodulin-binding protein of the nervous system. Proc Natl Acad Sci U S A. 1979;76(12):6270–3. doi: 10.1073/pnas.76.12.6270. PubMed PMID: 293720; PubMed Central PMCID: PMCPMC411845.

71. Liu J, Farmer JD, Jr., Lane WS, Friedman J, Weissman I, Schreiber SL. Calcineurin is a common target of cyclophilin-cyclosporin A and FKBP-FK506 complexes. Cell. 1991;66(4):807–15. doi: 10.1016/0092-8674(91)90124-h. PubMed PMID: 1715244.

72. Shirakata M, Friedman FK, Wei Q, Paterson BM. Dimerization specificity of myogenic helix-loop-helix DNA-binding factors directed by nonconserved hydrophilic residues. Genes Dev. 1993;7(12A):2456–70. doi: 10.1101/gad.7.12a.2456. PubMed PMID: 8253390.

73. Olson EN, Perry M, Schulz RA. Regulation of muscle differentiation by the MEF2 family of MADS box transcription factors. Dev Biol. 1995;172(1):2–14. doi: 10.1006/dbio.1995.0002. PubMed PMID: 7589800.

74. Au-Yeung N, Horvath CM. Transcriptional and chromatin regulation in interferon and innate antiviral gene expression. Cytokine Growth Factor Rev. 2018;44:11–7. Epub 20181022. doi: 10.1016/j.cytogfr.2018.10.003. PubMed PMID: 30509403; PubMed Central PMCID: PMCPMC6281172.

75. Martinez-Pena YVI, Akaaboune M. The Metabolic Stability of the Nicotinic Acetylcholine Receptor at the Neuromuscular Junction. Cells. 2021;10(2). Epub 20210209. doi: 10.3390/cells10020358. PubMed PMID: 33572348; PubMed Central PMCID: PMCPMC7916148.

76. Washington TA, Schrems ER. Skeletal Muscle Damage and Inflammation. Adv Exp Med Biol. 2025;1478:185–212. doi: 10.1007/978-3-031-88361-3_9. PubMed PMID: 40879941.

77. Koike TE, Fuziwara CS, Brum PC, Kimura ET, Rando TA, Miyabara EH. Muscle Stem Cell Function Is Impaired in beta2-Adrenoceptor Knockout Mice. Stem Cell Rev Rep. 2022;18(7):2431–43. Epub 20220304. doi: 10.1007/s12015-022-10334-y. PubMed PMID: 35244862.

78. Pearen MA, Ryall JG, Lynch GS, Muscat GE. Expression profiling of skeletal muscle following acute and chronic beta2-adrenergic stimulation: implications for hypertrophy, metabolism and circadian rhythm. BMC Genomics. 2009;10:448. Epub 20090923. doi: 10.1186/1471-2164-10-448. PubMed PMID: 19772666; PubMed Central PMCID: PMCPMC2758907.

79. Ryall JG, Church JE, Lynch GS. Novel role for ss-adrenergic signalling in skeletal muscle growth, development and regeneration. Clin Exp Pharmacol Physiol. 2010;37(3):397–401. Epub 20090928. doi: 10.1111/j.1440-1681.2009.05312.x. PubMed PMID: 19793099.

80. Endo T. Postnatal skeletal muscle myogenesis governed by signal transduction networks: MAPKs and PI3K-Akt control multiple steps. Biochem Biophys Res Commun. 2023;682:223–43. Epub 20230927. doi: 10.1016/j.bbrc.2023.09.048. PubMed PMID: 37826946.

81. Girardi F, Le Grand F. Wnt Signaling in Skeletal Muscle Development and Regeneration. Prog Mol Biol Transl Sci. 2018;153:157–79. Epub 20180108. doi: 10.1016/bs.pmbts.2017.11.026. PubMed PMID: 29389515.

82. Liu SY, Chen LK, Jhong YT, Chen CW, Hsiao LE, Ku HC, et al. Endothelin-1 impairs skeletal muscle myogenesis and development via ETB receptors and p38 MAPK signaling pathway. Clin Sci (Lond). 2024;138(12):711–23. doi: 10.1042/CS20240341. PubMed PMID: 38804865.

83. Bensalah M, Muraine L, Boulinguiez A, Giordani L, Albert V, Ythier V, et al. A negative feedback loop between fibroadipogenic progenitors and muscle fibres involving endothelin promotes human muscle fibrosis. J Cachexia Sarcopenia Muscle. 2022;13(3):1771–84. Epub 20220322. doi: 10.1002/jcsm.12974. PubMed PMID: 35319169; PubMed Central PMCID: PMCPMC9178170.

84. Dinenno FA, Joyner MJ. Alpha-adrenergic control of skeletal muscle circulation at rest and during exercise in aging humans. Microcirculation. 2006;13(4):329–41. doi: 10.1080/10739680600618843. PubMed PMID: 16611594.

85. Liu Y, Contreras M, Shen T, Randall WR, Schneider MF. Alpha-adrenergic signalling activates protein kinase D and causes nuclear efflux of the transcriptional repressor HDAC5 in cultured adult mouse soleus skeletal muscle fibres. J Physiol. 2009;587(Pt 5):1101–15. Epub 20090105. doi: 10.1113/jphysiol.2008.164566. PubMed PMID: 19124542; PubMed Central PMCID: PMCPMC2673778.

86. Iannotti FA, Silvestri C, Mazzarella E, Martella A, Calvigioni D, Piscitelli F, et al. The endocannabinoid 2-AG controls skeletal muscle cell differentiation via CB1 receptor-dependent inhibition of Kv7 channels. Proc Natl Acad Sci U S A. 2014;111(24):E2472–81. Epub 20140603. doi: 10.1073/pnas.1406728111. PubMed PMID: 24927567; PubMed Central PMCID: PMCPMC4066524.

87. Ono Y, Gnocchi VF, Zammit PS, Nagatomi R. Presenilin-1 acts via Id1 to regulate the function of muscle satellite cells in a gamma-secretase-independent manner. J Cell Sci. 2009;122(Pt 24):4427–38. Epub 20091117. doi: 10.1242/jcs.049742. PubMed PMID: 19920078; PubMed Central PMCID: PMCPMC2787457.

