## Supplemental Table 1 for "*Tceal7* is a BRG1-regulated target of calcineurin signaling that promotes myoblast differentiation"

Primers for qPCR (5’-3’)

| | |
|---|---|
| *Tceal7* | FW: ACTTGTGGCCAAGGAGAAGA |
| | REV : GGGTTGTTCATCCTCCCTCT |
| *CnA* | FW: GTTTAAACAAGAATGTAAAATAAA |
| | REV : GAATCGGTCTAATTTTCTGATGTC |
| *Myogenin* | FW: GTCCCAACCCAGGAGATCATTTGCTC |
| | REV : CCACTTAAAAGCCCCCTGCTA |
| *Eef1A1* | FW: GGCTTCACTGCTCAGGTGATTATC |
| | REV: ACACATGGGCTTGCCAGGGAC |

Primers for ChIP (5’-3’)

| | |
|---|---|
| *Tceal7* | FW: CCGCCCGAAAGGTCTATATT |
| | REV: TTTTACCTTTCCCTGATATCTTCC |
| *Myogenin* | FW: ACACCAACTGCTGGGTGCCA |
| | REV: GAATCACATGTAATCCACTGG |
